# Adherent and suspension baby hamster kidney cells have a different cytoskeleton and surface receptor repertoire

**DOI:** 10.1101/2021.01.25.428059

**Authors:** Veronika Dill, Florian Pfaff, Aline Zimmer, Martin Beer, Michael Eschbaumer

## Abstract

Animal cell culture, with single cells growing in suspension, ideally in a chemically defined environment, is a mainstay of biopharmaceutical production. The synthetic environment lacks exogenous growth factors and usually requires a time-consuming adaptation process to select cell clones that proliferate in suspension to high cell numbers. The molecular mechanisms that facilitate the adaptation and that take place inside the cell are largely unknown. Especially for cell lines that are used for virus antigen production such as baby hamster kidney (BHK) cells, the restriction of virus growth through the evolution of undesired cell characteristics is highly unwanted. The comparison between adherently growing BHK cells and suspension cells with different susceptibility to foot-and-mouth disease virus revealed differences in the expression of cellular receptors such as integrins and heparan sulfates, and in the organization of the actin cytoskeleton. Transcriptome analyses and growth kinetics demonstrated the diversity of BHK cell clones and confirmed the importance of well-characterized parental cell clones and mindful screening to make sure that essential cellular features do not get lost during adaptation.

## 1. Introduction

Animal cell culture technology was first used for the production of virus antigen in vaccine production, but more recently bacterial or yeast production platforms for recombinant proteins are changed over to mammalian cell systems as well (1, 2). Important in both contexts are baby hamster kidney (BHK) cells, an adherently growing, anchor-dependent mammalian cell line with a fibroblast growth pattern in standard cell culture conditions including foetal bovine serum (FBS) (3, 4). The adaptation of adherent BHK cells to suspension growth in serum-free media combined with large-scale culture conditions is a success story for the production of recombinant proteins (e.g. factor VIII) with high yields (1, 2). Furthermore, BHK cells are susceptible to a wide range of viruses, including the virus of foot-and-mouth disease (FMD) (5). To meet the increasing demand for FMD vaccines in many parts of the world, large quantities of antigen must be produced as cheaply as possible, which is achievable by increasing viable cell densities on the cellular side and decreasing cost through simplification of up- and downstream processing of the antigen on the production side (6). Animal-component-free media (ACFM) reduce the risk of contamination by adventitious agents and additionally simplify the process when no serum has to be supplemented (6, 7). However, the adaptation of cells to grow in suspension without serum and in ACFM is a time-consuming and gradual process (6, 8). BHK cells easily reach cell densities of 3.5×10^6^ cells/mL in suspension but the adaptation always carries the risk of evolving undesired cell properties in comparison to the starting population that can result in poor growth performance, insufficient cell specific yields or tumorigenic assets (5, 7). In turn, changes to the cells can lead to unexpected virus variants during the culture adaptation process. Especially for vaccine antigen production a change in virus characteristics and antigenicity is highly undesirable. BHK cells already differ in their susceptibility and their possibility to propagate certain virus strains of FMDV (9). Obviously, the change from anchorage-dependent fibroblastic growth via aggregation and formation of spheroids to cells growing independently in suspension will be accompanied by significant changes to the cell itself (4, 6). The loss of integrin signaling due to the interruption of extracellular matrix-cell contact leads to changes in membrane protein expression and in the organization of the actin cytoskeleton which is an important recipient of integrin-mediated signaling (6, 10). At the same time, these integrins play an important role for FMDV infection because they are the receptor for FMDV in the natural host and the primary receptor in cell culture (11).

A more profound understanding of the cellular differences between adherently growing BHK cells and cells in suspension will open up new possibilities for cell engineering and simplify the selection of suitable cell clones for vaccine antigen production.

In this study, adherently growing BHK cells and ACFM-adapted suspension BHK cells were examined with regard to the expression of cellular receptors such as integrins and heparan sulfates (HS), their ability to present proteins on the cell surface and the organization of the cellular actin skeleton. Transcriptome analyses and growth kinetics provide information about the diversity of the tested BHK cell clones.

## 2. Materials and Methods

### 2.1 Cell lines and cell culture

The principal cell lines used were two adherent BHK21 clone 13 lines (derived from American Type Culture Collection [ATCC] CCL-10, held as CCLV-RIE 164 and 179 in the Collection of Cell Lines in Veterinary Medicine, Friedrich-Loeffler-Institut, Greifswald, Germany; in short: BHK164 or BHK179) and the suspension cell lines BHK21C13-2P (BHK-2P) provided by the European Collection of Authenticated Cell Cultures (ECACC; 84111301) and BHK-InVitrus (BHK-InV, also referred to as “line #8” as in (12); Sigma-Aldrich, St. Louis, USA).

For growth kinetics, the following cell lines were used in addition: BHK-2P maintained either in Glasgow’s Minimal Essential Medium (GMEM) with 10% FBS (“line #2”) or adapted to the ACFM Cellvento™ BHK200 (Merck KGaA, Darmstadt, Germany) in two different processes (lines “#4” and “#5”), plus three other suspension cell lines: BHK21-C, “line #6”; BHK21-Hektor, “line #7” and production BHK, “line #9” (12). All these cell lines were provided by Merck KGaA.

The adherent Chinese hamster ovary (CHO) cell lines CHO-K1 (ATCC CCL-61, held as CCLV-RIE 134) and CHO677 (CRL 2244, held as CCLV-RIE 1524) as well as IB-RS-2 cells (Instituto Biológico-Rim Suíno-2, held as CCLV-RIE 103) and Madin-Darby bovine kidney cells (in short: MDBK, held as CCLV-RIE261) were used as controls in flow cytometry experiments.

Culture conditions were 37 °C, 5% CO_2_, and for suspension cells 320 rpm in TubeSpin bioreactors (TPP Techno Plastic Products AG, Trasadingen, Switzerland) at 80% relative humidity. All adherent cell lines were cultured in Minimum Essential Medium Eagle (MEM) supplemented with Hanks’ and Earle’s salts (Sigma-Aldrich) with 10% FBS. All suspension cell lines were adapted to grow in Cellvento™ BHK200 medium (Merck KGaA) except cell line #2, which was cultured as described above.

Measurements of cell density and viability were performed using trypan blue (Bio-Rad, Hercules, CA, USA) and an automatic cell counter (model TC20, Bio-Rad).

### 2.2 Growth curve analysis of suspension cells

Growth curve analyses were conducted as batch experiments, performed in duplicates, two times individually, with a seeding density of 0.6×10^6^ cells/mL. Viable cell density (VCD) and viability in percent were measured daily up to six days after cell inoculation.

The cell-specific calculations were performed according to the equations given by (13): Specific Growth Rate: µ (h^-1^)

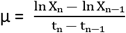

X: VCD (per mL); t: time points of sampling (hour); n and n-1: two consecutive sampling points.

Cell Division Number (cd):

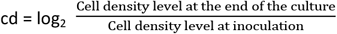

### 2.3 Flow Cytometry

Flow cytometric analyses were performed using the flow cytometry instrument FACSCanto II (BD Biosciences, Oxford, UK). Cell debris was gated out based on the forward and side scatter patterns. For actin staining, the median fluorescence intensity in the Alexa 488 channel was measured directly. For all antibody stains, the percentage of Alexa 488 or PE-positive cells was determined with an interval gate whose lower boundary was set based on an isotype control. All experiments were performed three times independently.

#### 2.3.1 Antibody staining of cellular surface receptors

BHK179, BHK-2P and BHK-InV cells were examined for the expression of integrin αvβ3, integrin αvβ8 and HS on the cellular surface. MDBK cells served as positive control for the expression of αvβ3, IB-RS-2 cells served as positive control for the expression of αvβ8 and CHO-K1 cells served as positive control for the expression of HS. CHO677 cells do not express any of the tested integrins nor HS and therefore served as negative control in all experiments. The following antibodies were used: for αvβ3, PE-conjugated clone LM609 (dilution 1:20) (mAb1976H, Merck KGaA); for αvβ8, primary antibody 11E8, clone 68, mouse IgG2a (dilution 1:100) (kindly provided by Dr. Stephen Nishimura) with secondary antibody goat anti-mouse IgG2a (dilution 1:1000) conjugated with Alexa Fluor 488 (Thermo Fisher Scientific); for HS, clone 10E4, mouse IgM (dilution 1:200) (Amsbio, Abingdon, UK) with secondary antibody goat anti-mouse IgM mu chain (dilution 1:2000) conjugated with Alexa Fluor 488 (ab150121, Abcam, Cambridge, UK). Only one primary antibody was used in each experiment.

Suspension cells were washed with phosphate-buffered saline (PBS). Adherent cells were detached using 10 mM EDTA in PBS and then washed with PBS. After washing, cells were filtered through a cell strainer (nylon mesh, 70 µm pore diameter, VWR, Radnor, USA) to remove cell clumps and were adjusted to 1×10^6^ cells/mL before 1 µL of LIVE/DEAD Fixable Violet dead cell stain (Thermo Fisher Scientific) was added to 1 mL cell suspension for 30 minutes. This step and all following steps were performed protected from light and at 4 °C. Primary antibody was incubated for 30 minutes and secondary antibody was incubated for 20-30 minutes. A washing step with 2 mL FACS buffer (PBS, 0.1% sodium azide, 0.1% BSA) and centrifugation at 300 × g, 5 min, 4 °C was performed after all steps. Finally, the stained cells were resuspended in 300 µL FACS buffer.

#### 2.3.2 Actin staining

Filamentous actin was stained using Phalloidin-iFluor 488 Reagent (ab176753, Abcam), prepared according to the datasheet. BHK179, BHK-2P and BHK-InV cells were detached as described above and fixed with 4% formaldehyde for 20 minutes at room temperature (RT). Cells were washed twice with PBS and 1×10^6^ fixed cells were permeabilized with 0.1 % Triton X-100 (Sigma-Aldrich) in PBS, followed by incubation at RT for 3 min. Cells were washed twice with PBS (centrifugation at 300 × g, 5 min) before adding the prepared phalloidin stock solution, diluted 1:1000 in FACS buffer and incubated for 20 min at RT. Lastly, cells were washed twice with FACS buffer and were resuspended in 300 µL FACS buffer. Stained cells were analyzed directly with a fluorescence microscope and used for FACS analysis.

#### 2.3.3 Transfection experiments

BHK164 and BHK-2P cells were transfected with either pEGFP-N1, a plasmid expressing enhanced green fluorescent protein (EGFP) (Clontech), or in another series of experiments with pVSV-G, a plasmid expressing the G protein of vesicular stomatitis Indiana virus (VSV), using Lipofectamine 3000 (Thermo Fisher Scientific) according to the manufacturer’s instructions. In short, 2 µg of plasmid DNA and 1.875 µL or 3.75 µL of lipofectamine diluted in Cellvento™ BHK200 medium were mixed and incubated at RT for 20 min to allow complex formation. Afterwards, these mixtures were transferred to cells in 12-well plates (approximately 4×10^5^ cells/mL) and medium was added up to 200 µL. The cells were incubated at 37 °C for 24 h.

For FACS analysis, the entire content of the wells was harvested after 24 h. The pEGFP-transfected cells were washed three times with PBS and were resuspend in 300 µL FACS buffer. One duplicate of pVSV-G-transfected cells was permeabilized with PBS with 0.1% (v/v) Triton™ X-100 for 5 min at RT prior to antibody staining, while the other remained non-permeabilized. Antibody staining was performed as described above with polyclonal rabbit serum VSV 274/E (dilution 1:500, kindly provided by Dr. Stefan Finke) and secondary antibody goat anti-rabbit IgG (H+L) conjugated with Alexa Fluor 488 (dilution 1:1000; Thermo Fisher Scientific).

### 2.4 Statistical Analysis

The data were analyzed using GraphPad Prism version 07.04 for Windows (GraphPad Software, La Jolla, USA). Ordinary one-way analysis of variance was done with Tukey’s post-hoc test to evaluate differences between treatment groups. The threshold for statistical significance was set at a p-value of 0.001.

### 2.5 Transcriptome analysis

#### 2.5.1 RNA extraction, library preparation and sequencing

Three independent batches of each of the adherent BHK179 cell line and the BHK-2P and BHK-InV suspension cell lines were used for transcriptome analysis. On average, 8.3×10^5^ BHK179 cells, 1.6×10^7^ BHK-2P cells and 1.4×10^7^ BHK-InV cells were lysed in TRIzol Reagent (Thermo Fisher Scientific). After adding 0.2 volumes of trichloromethane to the cell lysate, incubation and centrifugation, the aqueous phase was collected and mixed with one volume of 100% ethanol. From this, total RNA was extracted using the RNeasy Mini Kit (Qiagen, Hilden, Germany) with on-column DNase digestion with the RNase-Free DNase Set (Qiagen) following the manufacturer’s instructions. The ERCC RNA Spike-In Mix 1 (Thermo Fisher Scientific) was added and used as an internal control for all following steps. Next, polyadenylated mRNA was isolated from 1-3 µg of high-quality total RNA using the Dynabeads mRNA DIRECT Micro kit (Thermo Fisher Scientific). The mRNA was subsequently fragmented and used for cDNA synthesis and library preparation using the TruSeq Stranded mRNA LT Sample Prep Kit (Illumina, San Diego, USA) according to the manufacturer’s instructions. Sequencing was performed at the Heinrich-Pette-Institut, Hamburg, using NextSeq 500 (2×75 bp) and HiSeq 4000 (1×50 bp) equipment (Illumina).

#### 2.5.2 Statistical analysis of differential gene expression

A quality check of the raw reads from each sequencing library was performed using FastQC (version 0.11.9; Babraham Institute, Cambridge, UK) with emphasis on the read length distribution and adapter contamination before and after adapter trimming using Trim Galore (version 0.6.4_dev, Babraham Institute) and Cutadapt (version 2.10, (14)). The trimmed raw reads were then quasi-mapped to a public reference of *Mesocricetus auratus* using Salmon (15). In short, the genomic and transcript references of GCF_000349665.1_MesAur1.0 were received from the National Center for Biotechnology Information (NCBI) and combined with reference sequences of the ERCC spike-in controls. A decoy-aware transcriptome index (16) was constructed in order to allow selective alignment with Salmon. For quantification, ten bootstrap replicates were used and compensation for sequence-specific biases, GC biases and 5’ or 3’ positional bias was enabled.

Subsequently, the quantification data for the ERCC spike-in controls was correlated to their theoretical concentration, in order to detect problems during sample preparation.

Furthermore, the gene-specific quantification data were initially filtered (only genes that appear in more than one replicate with over 15 counts), normalized using regularized log transformation (rlog), and used for explorative data analysis, e.g. with PCA. Differentially expressed genes were identified and subsequently used for pathway analysis using DESeq2 (17). The expression data were analyzed for significantly differently expressed genes between the cell line pairs BHK-2P and BHK179, BHK-InV and BHK-2P, and BHK-InV and BHK179. Subsequently, the resulting binary logarithmic (log_2_) fold change was adjusted using functions “lfcShrink” and “apglm” implemented in DESeq2 in order to compensate for low or variable read counts that may lead to variance overestimation (18).

The criteria for a significantly differently expressed gene were set to a maximum adjusted p-value of 0.05 and a minimal absolute value of the shrunk log_2_ fold change of 1. Pathway and enrichment analysis was conducted using the function “enrichGO” in clusterProfiler (version 3.16.0).

#### 2.5.3 Data Availability

The raw sequencing data along with deduced Salmon read count tables and substantial metadata are available at ArrayExpress (http://www.ebi.ac.uk/arrayexpress) under the accession number E-MTAB-XXXX.

## 3. Results

### 3.1 The susceptibility of a cell line for FMDV is independent of its individual growth rate and cell division number

Batch experiments with nine different BHK suspension cell lines were conducted over a period of six days and viable cell density (VCD) and viability were measured daily. Based on previous observations that the cell lines differed in their susceptibility for FMDV serotype Asia-1 (12), it was examined whether an elevated cellular metabolism and growth rate is linked to a reduced susceptibility for infection with this virus (Table 1). Two of three susceptible BHK cell lines are slowly growing cells with small cell division numbers (cd) of 0.60 (#3) and 0.70 (#9), while the third susceptible cell line has the highest growth rate and cd (#8, µ=0.035, cd=1.62) among all tested cell lines. The withdrawal of serum and adaptation to grow in ACFM seems to play a minor role in our set of examined cell lines. Cell line #2 is the slowly growing, serum-dependent precursor of cell line #1. During adaptation, cells were selected for high VCD, linked to a high growth rate and cd. Cell lines #4 and #5 are also derived from cell line #2 but differ in the mode of adaptation to ACFM. Susceptibility for FMDV serotype Asia-1 was not acquired during adaptation, probably because the original cell line had already been resistant. The opposite was seen for cell line #3. When growing adherently, this cell line was susceptible to FMDV Asia-1 and kept this characteristic during adaptation to grow in ACFM, but also did not acquire the rapid growth profile of other suspension cells.

**Table 1.**
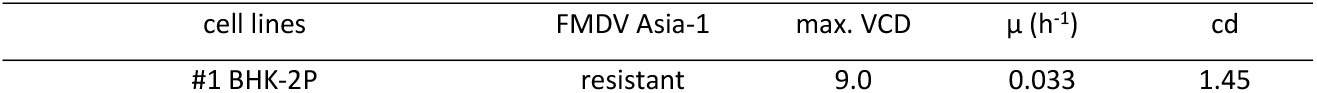
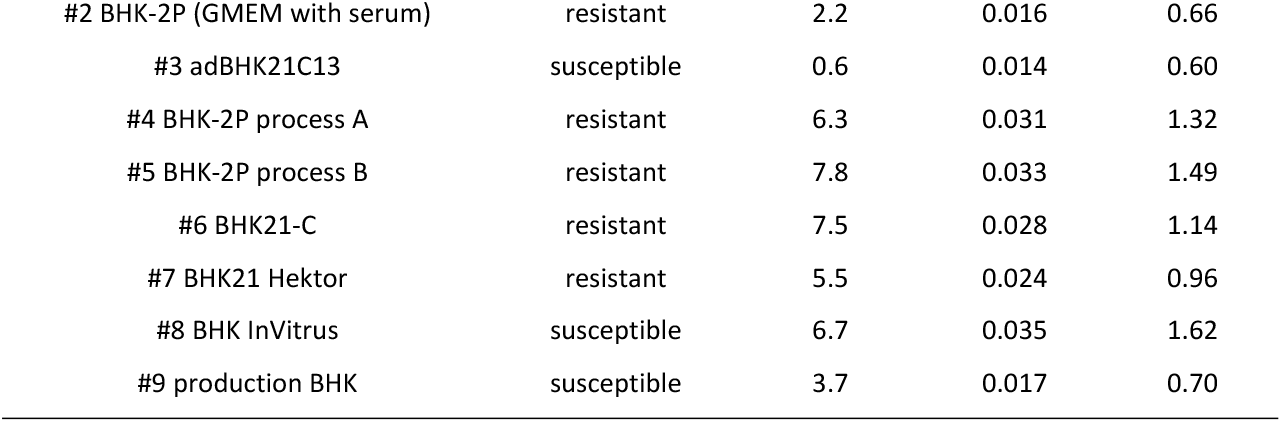
Maximum viable cell density (VCD), growth rate (µ (h^-1^)) and cell division number (cd) of different BHK cell lines in conjunction with the susceptibility for infection with FMDV Asia-1.

### 3.2 Suspension cells present less primary receptors but more secondary receptors for FMDV on their surface than adherent BHK cells

The first step for a virus to successfully infect a cell is the binding of the viral particle to receptor molecules on the cell surface. To analyze the differences between adherently growing BHK cells and BHK cells in suspension, antibodies to cellular receptors involved in the extracellular matrix interaction and known to play an important role for FMDV infection were tested. Integrins αvβ3 and αvβ8 were examined as representative cell receptors that recognize the RGD (Arg-Gly-Asp) binding motif and interact with serum components and the extracellular matrix but are also used by FMDV as its natural primary receptors. Neither of the tested suspension cell lines (BHK-InV and BHK-2P) expressed integrin αvβ3 or αvβ8 on the cell surface. While the difference between adherent BHK and suspension BHK was not significant for integrin αvβ8 (Figure 1b), a significantly higher percentage of adherent BHK179 expressed integrin αvβ3 compared to the suspension cell lines BHK-InV and BHK-2P (Figure 1a).

**Fig 1.**
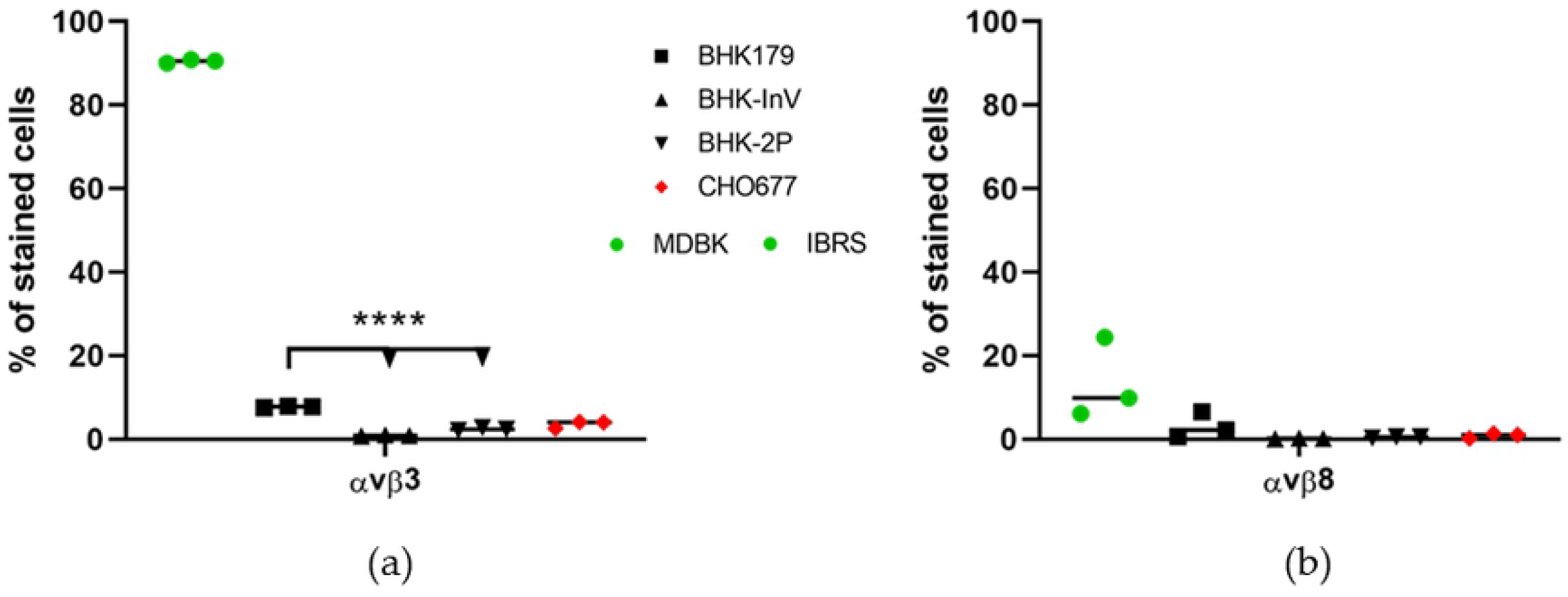
Expression of integrin αvβ3 and αvβ8 on the cellular surface of the adherently growing cell line BHK179 and two suspension cell lines BHK-InV and BHK-2P. Cells were stained with antibodies targeting αvβ3 (a) or αvβ8 (b) and were analyzed by flow cytometry. MDBK cells served as positive control for integrin αvβ3, while IB-RS-2 (IBRS) cells served as positive control for the expression of αvβ8 (green dots). BHK179 cells are presented as black squares, while the suspension cell lines are represented as triangles. The negative control cell line CHO677 is shown as red diamonds. Experiments were performed three times independently. Significance code: **** p < 0.0001.

High amounts of HS were presented on the surface of both suspension cell lines. The expression of HS was highly divergent between the replicate cultures of adherent BHK179 cells. Compared to the suspension cell lines, fewer cells in the BHK179 cultures were HS-positive, even though the difference was not significant at the predetermined level of significance (Figure 2).

**Fig 2.**
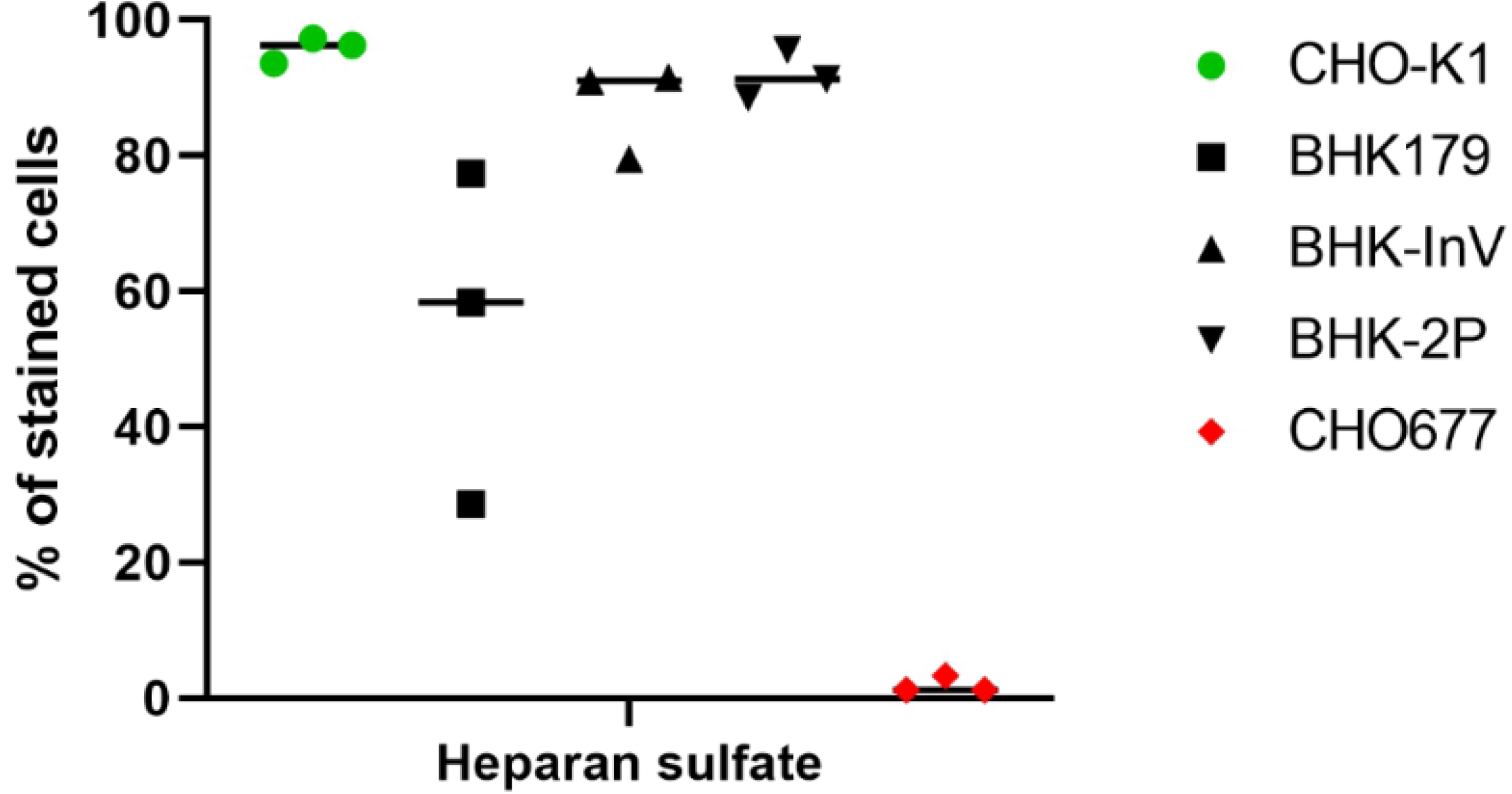
Expression of heparan sulfate (HS) on the cellular surface of the adherently growing cell line BHK179 and the two suspension cell lines BHK-InV and BHK-2P. Cells were incubated with an antibody targeting HS and were analyzed by flow cytometry. CHO-K1 cells served as positive control (green circles), while CHO677 cells served as negative control (red diamonds). BHK179 cells are shown as black squares and the suspension cell lines are represented by triangles. Experiments were performed three times independently.

Staining and microscopical analysis revealed large differences in the arrangement of the actin cytoskeleton between adherent and suspension cells (Figure 3). The actin filaments in the adherent BHK179 cells are uniformly distributed over the entire cell in a fine reticular pattern (Figure 3a), whereas cells growing in suspension had diffuse actin staining (Figure 3b). Interestingly, the quantity of filamentous actin was different between the suspension cell lines BHK-InV and BHK-2P. The median fluorescent intensity (MFI) after phalloidin staining was significantly higher in BHK-InV cells compared to BHK-2P, which indicates a higher filamentous actin content in these cells (Figure 3c).

**Fig 3.**
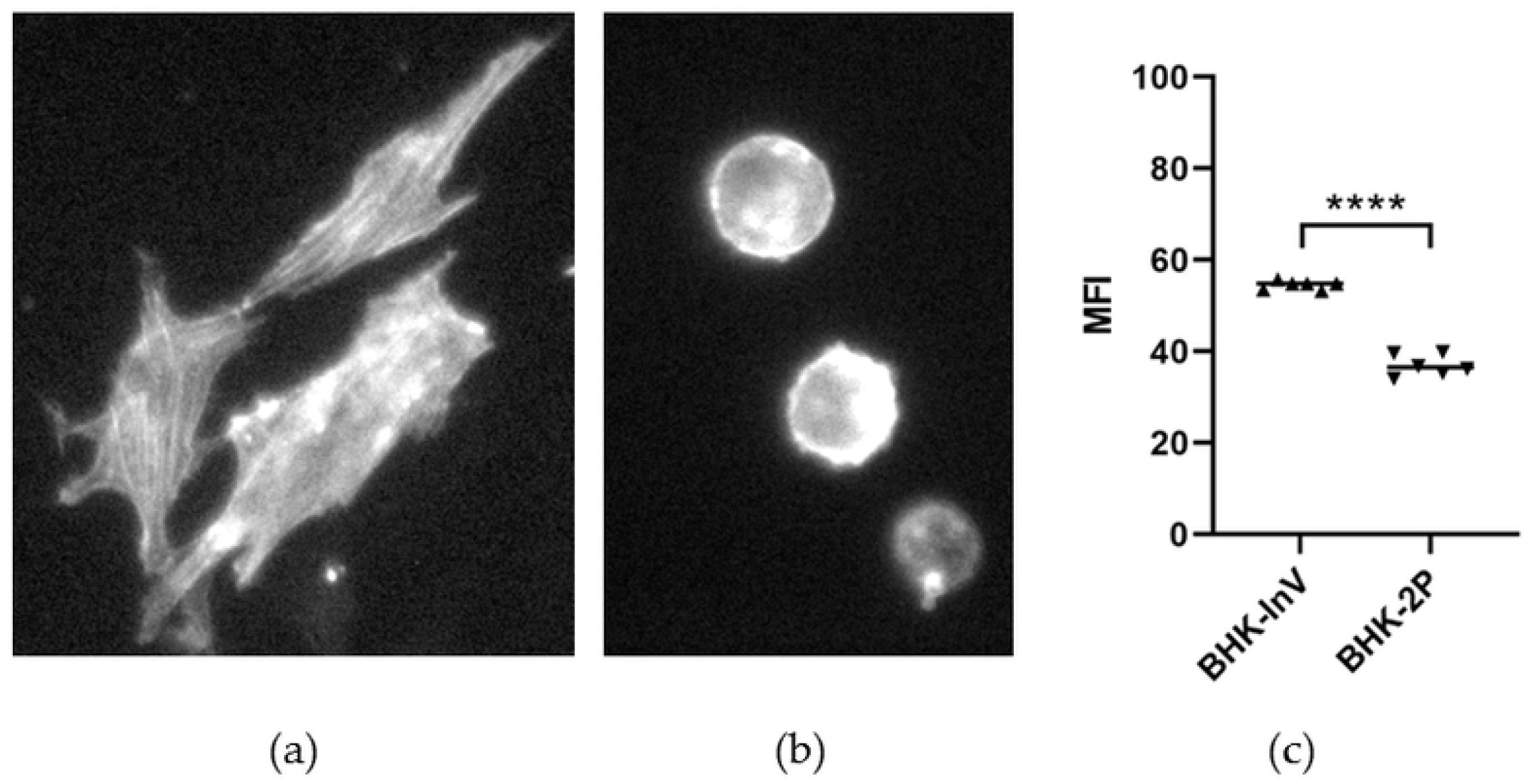
Quantity and distribution of filamentous actin. The microscopy images give examples of the distribution of filamentous actin in BHK179 cells in adherent growth conditions (a) and in one of the suspension cell lines (BHK-2P) (b). Microscopically, there was no difference between the two suspension cell lines. The median fluorescent intensity (MFI) of filamentous actin staining in BHK-InV and BHK-2P (c) was measured by flow cytometry in two replicates per experiment. The experiment has been performed three times independently. Significance code: **** p < 0.0001.

### 3.3 Suspension cells differ in transfection efficiency but not in their ability to display surface proteins

Because of the differences in actin conformation between adherent and suspension cells, the cells were examined with regard to their transfectability and the ability to present proteins on their surface. In general, the transfection efficacy, measured by the expression of EGFP, is reduced in suspension cells (Figure 4a). When using a low dose of transfection reagent, a significantly higher percentage of adherent cells expressed EGFP compared to the suspension cells. When a high dose of transfection reagent was used, a higher percentage of suspension cells were successfully transfected, although the difference between high and low doses was not significant. To answer the question whether suspension cells are still able to present proteins such as cellular receptors on their surface, cells were transfected with a plasmid coding for the VSV G protein (Figure 4b). One series of cells was permeabilized prior to staining to account for the possibility that the G protein was expressed but not displayed on the cellular surface. There was no significant difference in the ability to express and present the VSV G protein between adherent and suspension cells, neither when using a low or a high dose of transfection reagent.

**Fig 4.**
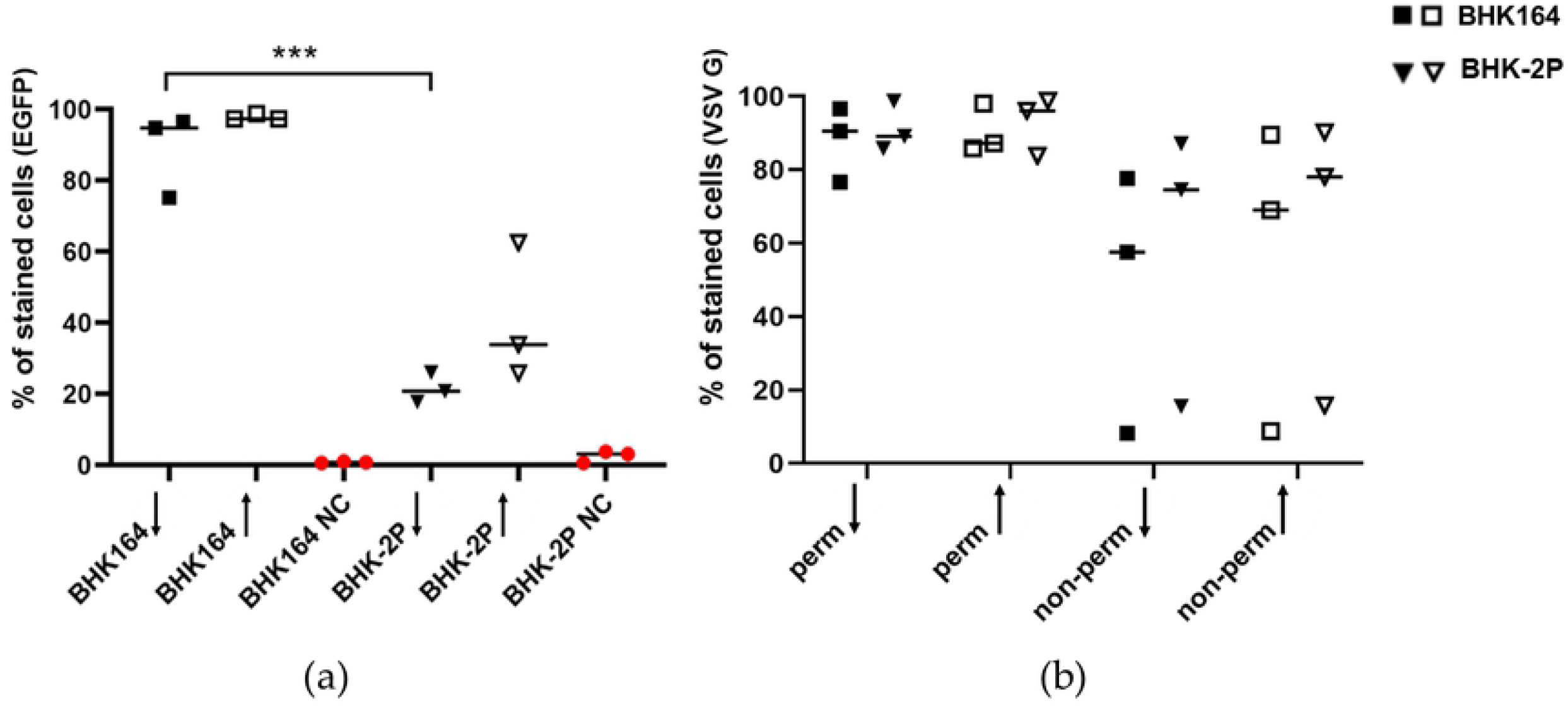
Transfection efficacy and ability to display proteins on the cellular surface of adherent and suspension cells. Adherent BHK164 cells and BHK-2P suspension cells were transfected with plasmids expressing EGFP (a) or VSV G protein (b). For transfection, either a low dose (downward arrow) or a high dose (upward arrow) of transfection reagent was used. The percentage of successfully transfected cells was measured with flow cytometry after 24 h incubation at 37 °C. The experiments were performed three times independently. Significance code: *** p < 0.001.

### 3.4 Transcriptome analysis reveals differences in the expression of genes related to the extracellular matrix between suspension cells

A transcriptome analysis was performed to further investigate differences between adherently growing BHK cells and BHK cells in suspension and to elucidate why suspension cells differ in their susceptibility for FMDV. An adherent BHK cell line (BHK179) cultured in MEM supplemented with FBS as well as a fast-growing FMDV-susceptible cell line (BHK-InV) and a fast-growing resistant cell line (BHK-2P), both adapted to ACFM, were chosen for the analysis.

A principal component analysis (PCA) highlighted the close relationship between the biological replicates of the individual cell lines (Figure 5a). All replicates of the same cell line are located closely together in the plot with little indication of batch-to-batch effects. The first principal component (accounting for 81% of variance in the normalized dataset) was interpreted as the transcriptional difference between cells growing in suspension and cells growing in adherence, while the second principal component (15% of variance) represented the difference between the suspension cell lines BHK-2P and BHK-InV, possibly associated with susceptibility to FMDV infection. Together, the first two principal components were able to explain 96% of the variance found within the dataset, which indicates that the transcriptome analysis identified relevant differences between the three cell lines. The smallest number of differentially expressed genes was found when comparing BHK-InV with BHK-2P (590 up-regulated; 304 down-regulated). Similar numbers were seen in the comparison of either BHK-2P or BHK-InV individually with BHK179 (1527 and 1519 up-regulated; 1308 and 943 down-regulated, respectively) (Figure 5b). When the two suspension cell lines (BHK-InV and BHK-2P) were combined and compared to the adherent BHK179 cell line, 1814 genes were differently expressed. While 411 genes are exclusively differently expressed when comparing BHK179 with BHK-InV, nearly twice as many (701 genes) are exclusively differently expressed between BHK179 and BHK-2P. Between the two suspension cell lines, only 104 genes were exclusively differently expressed.

**Fig 5.**
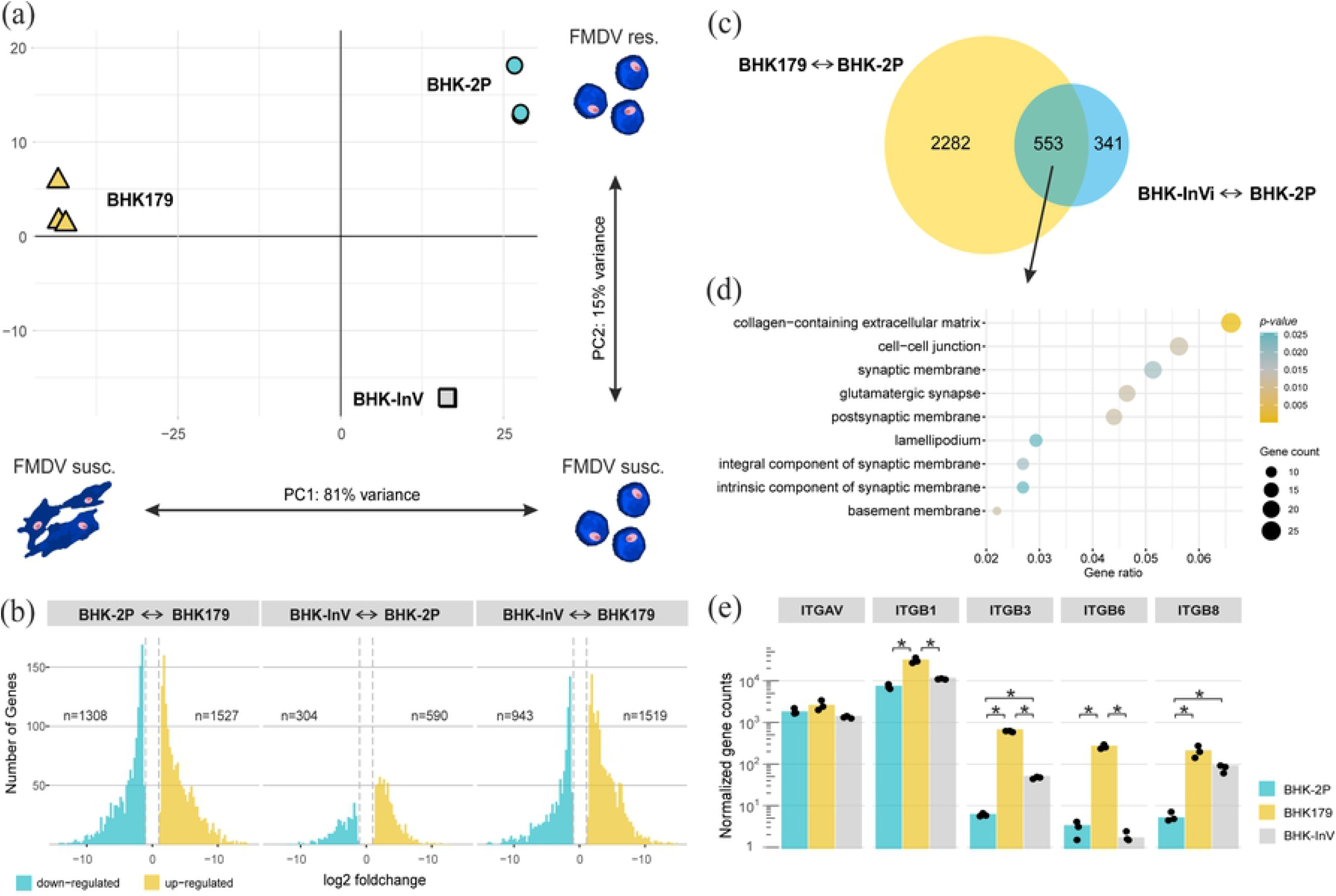
Explorative and differential gene expression analysis between adherent and suspension cell lines. (a) The principal component analysis (PCA) visualizes the difference in gene expression patterns between the adherent BHK179 cells (yellow triangles) and the two suspension cell lines BHK-InV (grey squares) and BHK-2P (blue circles). (b) The number and fold change of differentially expressed genes for each of the three possible comparisons. Down- and up-regulated genes are indicated in blue and yellow, respectively. Dashed lines indicate the cutoff value of |log2 foldchange| > 1. (c) The Venn diagram summarizes differentially expressed genes when comparing FMDV-susceptible cell lines BHK179 and BHK-InV with the FMDV-resistant cell line BHK-2P. (d) The subset of 553 genes from (c) was used for GO term enrichment analysis focusing on cellular components. (e) Normalized gene counts for different integrin genes in the analyzed cell lines BHK179, BHK-2P and BHK-InV. The integrin genes ITGAV, ITGB1, ITGB3, ITGB6 and ITGB8, corresponding to the integrin subunits αv, β1, β3, β6 and β8, respectively, were analyzed. Significance code: * p < 0.05.

When comparing the set of differentially expressed genes of the FMDV-susceptible cell lines BHK179 and BHK-InV with the FMDV-resistant cell line BHK-2P, a subset of 553 genes were found to be differently expressed in both comparisons (Figure 5c). As these genes might be involved in FMDV susceptibility, a GO term enrichment analysis was conducted and revealed that many of these genes are related to the cellular components of the extracellular matrix (ECM), cell membranes and cell-cell junctions (Figure 5d).

Since we observed differences between the cell lines related to ECM interactions, the transcriptome was reanalyzed with a focus on the expression of integrins αvβ1, αvβ3, αvβ6, αvβ8, all potentially relevant for susceptibility to FMDV infection (Figure 5e). All three cell lines expressed similar quantities of ITGAV transcripts, coding for integrin subunit αv but showed differential expression for the integrin β subunits. In detail, the suspension cell lines BHK-2P and BHK-InV expressed significantly less ITGB1, ITGB3 and ITGB6 in comparison to the adherent cell line BHK179. ITGB3 and ITGB8 were significantly differentially expressed between the suspension cell lines BHK-2P and BHK-InV. Interestingly, ITGB8 was the only integrin subunit coding gene that was highly overexpressed in both FMDV-susceptible cell lines BHK179 and BHK-InV when compared to the FMDV-resistant cell line BHK-2P (Figure 5e).

## 4. Discussion

The adaptation of cells to grow in suspension and in serum-free media is a time-consuming and gradual but necessary process in biotechnology to generate high-yielding cell lines used to produce antibodies, vaccine antigen or other proteins (1, 8). Indeed, this process is always fraught with the risk that cells acquire undesired properties compared to the original cell material (7). In the case of vaccine antigen production the susceptibility of the cell line for the target virus is of major importance. For foot-and-mouth disease virus (FMDV) it is a well know phenomenon that certain BHK cell lines are resistant to infection or lose their susceptibility during repeated subculturing (19-21). In addition, the antigenic properties of the virus can be variable depending on the host cell (9). Suspension cells grow independently of a surface and are engulfed by oxygen-rich nutrient media, allowing them to grow quickly. The rapid multiplication of cells is a required feature to scale up the culture to large volumes easily and in short time. The removal of serum from the medium is necessary to reduce costs, the risk of contamination and to simplify downscale processes (22, 23).

Batch growth analysis of different BHK suspension cell lines was intended to determine if the selection towards a fast-growing cell population in serum-free conditions is detrimental to the propagation of FMDV. In fact, two of three susceptible cell lines did grow slowly but their susceptibility to FMDV was found to be independent of their growth metabolism. It is more likely that FMDV susceptibility is either lost or retained during the initial adaptation to suspension growth.

The detachment of the cells from the surface and the withdrawal of serum in the following steps of adaptation to suspension growth leads to changes in the expression level of surface proteins and proteins involved in the cytoskeleton (7).

Integrins are an important family of cell surface receptors that are integrated into the cell membrane and mediate cell-to-cell signaling as well as a strong connection between the cell and the surface through the binding of extracellular matrix (ECM) components (10, 24). Cell cycle regulation, organization of the cytoskeleton and the translocation of nascent receptor molecules to the cell membrane depend on integrins (24). For many integrins, the binding of the ECM as well as the binding of ligands provided to the cells through the addition of serum to the culture medium is mediated by the RGD motif (6, 24) and it is likely to change when the cell switches from adherent to suspension growth. The recognition of the RGD motif by integrins is also used by FMDV to initiate a receptor-mediated endocytosis of the virus particle (25). The RGD motif in VP1 of FMDV is highly conserved (26), and especially the integrins αvβ1, αvβ3, αvβ6 and αvβ8 are known receptors used by FMDV in vivo and in vitro (27-30). Because the suspected differences in the cell surface proteome between adherent and suspension cells may also be crucial for the differences in susceptibility to FMDV infection, antibodies binding to the integrin αvβ3 and αvβ8 were used to stain an adherent BHK cell line as well as a susceptible and a resistant suspension BHK cell line. Notwithstanding the high level of ITGB8 mRNA expression in BHK179 and BHK-InV cells, αvβ8 integrin does not appear to be presented on the surface of BHK cells in general. Integrin αvβ3 was found to a minor degree on the adherent cell line, but not on the suspension cell lines.

Apart from integrins, ligand binding, internalization events and intracellular signaling can be mediated by heparan sulfate (HS) proteoglycans (31) and the acquisition of HS as a secondary receptor is often observed after repeated passaging of FMDV in culture (32). Specific antibody staining revealed abundant HS on both suspension cell lines and to a lesser extent also on the adherent BHK cells. Mutations in the genome of FMDV, leading to changes in the viral capsid proteins to allow the usage of HS as receptor, are known to attenuate the virus in the natural host (33). The availability of HS might therefore be a crucial factor for changed antigenicity of FMDV in dependence of the cell line the virus has been adapted to.

Because cellular detachment is linked to changes in the actin-myosin contractile mechanism (7) and culturing cells in suspension brings different requirements for survival, e.g. resistance to high shear stress due to agitation of the culture fluid (6), the actin skeleton has been of major interest in this study. Microscopical analysis revealed a conformational rearrangement of the cytoskeleton from a reticular distribution in adherent growing cells to a dense spherical structure in suspension conditions. This might be an adjustment mechanism to the above-mentioned high shear stress and is also known to occur in CHO cells in the same conditions (6). But it is also noteworthy that while there was no significant difference in the expression of the actin genes between BHK-InV and BHK-2P cells at the mRNA level (data not shown), significantly more filamentous actin protein was detected in the FMDV-susceptible BHK-InV than in the FMDV-resistant BHK-2P. This may be linked to different polymerization rates of globular actin into filamentous actin. At least in adherent cells the entry of FMDV depends on actin dynamics. Viral entry induces actin ruffles and a disruption of the actin filament network through the conversion of filamentous actin to globular actin has been described in the early stage of infection (34-36). Judging from to the cytopathic effect (CPE) seen in adherent BHK cells when infected with FMDV the cytoskeleton goes through intensive rearrangements in many of its components (36). Such CPE is not seen in suspension cells due to their already spherical shape but the elevated actin content may give some advantage to the virus. Because of the differences in the actin cytoskeleton, transfection experiments were conducted to compare the transfection efficiency between suspension and adherent BHK and to determine if the dense cytoskeleton of the suspension cells hinders the presentation of proteins on the cellular surface. It is common knowledge that suspension cell lines are very difficult to transfect, which is caused by reduced attachment of the transfection complex to the cell membrane and subsequently reduced uptake of the DNA payload (37). This was also confirmed by our experiments. But although the transfection efficiency was reduced in BHK-2P cells compared to the adherent BHK cell line, there was no difference in the ability to display the transfected VSV G protein on the cellular surface. Lastly, transcriptome analysis was performed to give more information about the transcriptional differences between adherent and suspension BHK cells on the one hand and between FMDV-susceptible and FMDV-resistant BHK cell lines on the other hand. Our results are in line with other studies of transcriptomic changes in CHO, MDCK or HEK cells during the changeover from adherent to suspension growth. In general, most changes are related to cytoskeletal structure, cellular metabolism and ECM component interactions and are upregulated for suspension cells (7, 38, 39). Surprisingly, the FMDV-resistant BHK-2P suspension cell showed downregulation of these gene sets, which might also explain the reduced actin content in BHK-2P compared to the BHK-InV cells. Specific analysis of integrin subunit expression revealed similar levels of integrin subunit αV transcripts in all three cell lines. The highest expression in all cell lines was found for the integrin subunit β1. It is important to note that integrin β1 can form heterodimers with ten different α chains and is constitutively expressed by most mammalian cells (6, 40). Similarly, the αV subunit is able to form a heterodimer with five different β chains, but β6 and β8 are *only* presented on the cell surface in combination with αV (40). This is accurately reflected by the observed abundance of transcripts of the single integrin subunits. The elevated transcription of β3 in adherent cells also correlates well with the higher signal for integrin αvβ3 in the antibody staining experiments. The suspension cell lines BHK-2P and BHK-InV expressed significantly less ITGB1, ITGB3 and ITGB6 mRNA in comparison to the adherent cell line.

## 5. Conclusions

In conclusion, it is of critical importance in the preparation of suspension cells to use a parental cell clone with well-known characteristics and to screen the resultant cell population carefully during the adaptation process to ensure that the cell line retains essential features, in this case the susceptibility to FMDV. In addition, modern methods such as FACS analysis and transcriptomics should be used for further characterization of production cell lines.

## Author Contributions

conceptualization, VD and ME; methodology, VD, ME, FP; formal analysis, VD and FP; investigation, VD; resources, AZ; writing—original draft preparation, VD; writing—review and editing, VD, ME, AZ, MB, FP; visualization, VD, FP, ME; supervision, ME; project administration, MB; funding acquisition, MB, AZ.

## Acknowledgments

We thank Patrick Zitzow for his excellent technical support.

## References

1. Merten OW. Advances in cell culture: anchorage dependence. Philos Trans R Soc Lond B Biol Sci. 2015;370(1661):20140040.

2. Dingermann T. Recombinant therapeutic proteins: production platforms and challenges. Biotechnol J. 2008;3(1):90–7.

3. Hernandez R, Brown DT. Growth and maintenance of baby hamster kidney (BHK) cells. Curr Protoc Microbiol. 2010;Chapter 4:Appendix 4H.

4. Cruz HJ, Moreira JL, Stacey G, Dias EM, Hayes K, Looby D, et al. Adaptation of BHK cells producing a recombinant protein to serum-free media and protein-free medium. Cytotechnology. 1998;26(1):59–64.

5. Guskey LE, Jenkin HM. Adaptation of BHK-21 cells to growth in shaker culture and subsequent challenge by Japanese encephalitis virus. Appl Microbiol. 1975;30(3):433–8.

6. Walther CG, Whitfield R, James DC. Importance of Interaction between Integrin and Actin Cytoskeleton in Suspension Adaptation of CHO cells. Appl Biochem Biotechnol. 2016;178(7):1286–302.

7. Kluge S, Benndorf D, Genzel Y, Scharfenberg K, Rapp E, Reichl U. Monitoring changes in proteome during stepwise adaptation of a MDCK cell line from adherence to growth in suspension. Vaccine. 2015;33(35):4269–80.

8. Costa AR, Withers J, Rodrigues ME, McLoughlin N, Henriques M, Oliveira R, et al. The impact of cell adaptation to serum-free conditions on the glycosylation profile of a monoclonal antibody produced by Chinese hamster ovary cells. N Biotechnol. 2013;30(5):563–72.

9. Bolwell C, Brown AL, Barnett PV, Campbell RO, Clarke BE, Parry NR, et al. Host cell selection of antigenic variants of foot-and-mouth disease virus. J Gen Virol. 1989;70 (Pt 1):45–57.

10. Wiesner S, Legate KR, Fassler R. Integrin-actin interactions. Cell Mol Life Sci. 2005;62(10):1081–99.

11. O’Donnell V, Pacheco JM, Gregg D, Baxt B. Analysis of foot-and-mouth disease virus integrin receptor expression in tissues from naive and infected cattle. J Comp Pathol. 2009;141(2-3):98–112.

12. Dill V, Hoffmann B, Zimmer A, Beer M, Eschbaumer M. Adaption of FMDV Asia-1 to Suspension Culture: Cell Resistance Is Overcome by Virus Capsid Alterations. Viruses. 2017;9(8).

13. Rourou S, van der Ark A, van der Velden T, Kallel H. A microcarrier cell culture process for propagating rabies virus in Vero cells grown in a stirred bioreactor under fully animal component free conditions. Vaccine. 2007;25(19):3879–89.

14. Martin M. Cutadapt removes adapter sequences from high-throughput sequencing reads. EMBnetjournal. 2011;17:10–2.

15. Patro R, Duggal G, Love MI, Irizarry RA, Kingsford C. Salmon provides fast and bias-aware quantification of transcript expression. Nat Methods. 2017;14(4):417–9.

16. Srivastava A, Malik L, Sarkar H, Zakeri M, Almodaresi F, Soneson C, et al. Alignment and mapping methodology influence transcript abundance estimation. Genome Biol. 2020;21(1):239.

17. Love MI, Huber W, Anders S. Moderated estimation of fold change and dispersion for RNA-seq data with DESeq2. Genome Biol. 2014;15(12):550.

18. Zhu A, Ibrahim JG, Love MI. Heavy-tailed prior distributions for sequence count data: removing the noise and preserving large differences. Bioinformatics. 2019;35(12):2084–92.

19. Clarke JB, Spier RE. Variation in the susceptibility of BHK populations and cloned cell lines to three strains of foot-and-mouth disease virus. Arch Virol. 1980;63(1):1–9.

20. Clarke JB, Spier RE. An investigation into causes of resistance of a cloned line of BHK cells to a strain of foot-and-mouth disease virus. Vet Microbiol. 1983;8(3):259–70.

21. Syusyukin AA, Tsvetkova NE, Kudryatseva GA, Syusyukin MS, Ni E. Cultures of foot-and-mouth disease virus in different sublines of BHK 21 cells. Veterinariya (Moscow). 1976;51:46–8.

22. Genzel Y, Fischer M, Reichl U. Serum-free influenza virus production avoiding washing steps and medium exchange in large-scale microcarrier culture. Vaccine. 2006;24(16):3261–72.

23. Merten OW. Development of serum-free media for cell growth and production of viruses/viral vaccines--safety issues of animal products used in serum-free media. Dev Biol (Basel). 2002;111:233–57.

24. Giancotti FG, Ruoslahti E. Integrin signaling. Science. 1999;285(5430):1028–32.

25. Stewart PL, Nemerow GR. Cell integrins: commonly used receptors for diverse viral pathogens. Trends Microbiol. 2007;15(11):500–7.

26. Carrillo C, Tulman ER, Delhon G, Lu Z, Carreno A, Vagnozzi A, et al. Comparative genomics of foot-and-mouth disease virus. J Virol. 2005;79(10):6487–504.

27. Jackson T, Clark S, Berryman S, Burman A, Cambier S, Mu D, et al. Integrin alphavbeta8 functions as a receptor for foot-and-mouth disease virus: role of the beta-chain cytodomain in integrin-mediated infection. J Virol. 2004;78(9):4533–40.

28. Jackson T, Mould AP, Sheppard D, King AM. Integrin alphavbeta1 is a receptor for foot-and-mouth disease virus. J Virol. 2002;76(3):935–41.

29. Jackson T, Sheppard D, Denyer M, Blakemore W, King AM. The epithelial integrin alphavbeta6 is a receptor for foot-and-mouth disease virus. J Virol. 2000;74(11):4949–56.

30. Jackson T, Sharma A, Ghazaleh RA, Blakemore WE, Ellard FM, Simmons DL, et al. Arginine-glycine-aspartic acid-specific binding by foot-and-mouth disease viruses to the purified integrin alpha(v)beta3 in vitro. J Virol. 1997;71(11):8357–61.

31. Bernfield M, Gotte M, Park PW, Reizes O, Fitzgerald ML, Lincecum J, et al. Functions of cell surface heparan sulfate proteoglycans. Annu Rev Biochem. 1999;68:729–77.

32. Jackson T, Ellard FM, Ghazaleh RA, Brookes SM, Blakemore WE, Corteyn AH, et al. Efficient infection of cells in culture by type O foot-and-mouth disease virus requires binding to cell surface heparan sulfate. J Virol. 1996;70(8):5282–7.

33. Sa-Carvalho D, Rieder E, Baxt B, Rodarte R, Tanuri A, Mason PW. Tissue culture adaptation of foot-and-mouth disease virus selects viruses that bind to heparin and are attenuated in cattle. J Virol. 1997;71(7):5115–23.

34. Han SC, Guo HC, Sun SQ, Jin Y, Wei YQ, Feng X, et al. Productive Entry of Foot-and-Mouth Disease Virus via Macropinocytosis Independent of Phosphatidylinositol 3-Kinase. Sci Rep. 2016;6:19294.

35. Armer H, Moffat K, Wileman T, Belsham GJ, Jackson T, Duprex WP, et al. Foot-and-mouth disease virus, but not bovine enterovirus, targets the host cell cytoskeleton via the nonstructural protein 3Cpro. J Virol. 2008;82(21):10556–66.

36. Bedows E, Rao KM, Welsh MJ. Fate of microfilaments in vero cells infected with measles virus and herpes simplex virus type 1. Mol Cell Biol. 1983;3(4):712–9.

37. Basiouni S, Fuhrmann H, Schumann J. High-efficiency transfection of suspension cell lines. Biotechniques. 2012;53(2).

38. Zhao L, Fu HY, Raju R, Vishwanathan N, Hu WS. Unveiling gene trait relationship by cross-platform meta-analysis on Chinese hamster ovary cell transcriptome. Biotechnol Bioeng. 2017;114(7):1583–92.

39. Malm M, Saghaleyni R, Lundqvist M, Giudici M, Chotteau V, Field R, et al. Evolution from adherent to suspension: systems biology of HEK293 cell line development. Sci Rep. 2020;10(1):18996.

40. Kumar CC. Signaling by integrin receptors. Oncogene. 1998;17(11 Reviews):1365–73.

